# ChatSpatial: Schema-Enforced Agentic Orchestration for Reproducible and Cross-Platform Spatial Transcriptomics

**DOI:** 10.64898/2026.02.26.708361

**Authors:** Chen Yang, Xianyang Zhang, Jun Chen

## Abstract

Spatial transcriptomics analyses often require coordinating specialized Python and R methods. When analytical intent is translated through ad hoc scripts or unconstrained LLM generated code, workflows can be difficult to inspect, rerun, or compare. We present ChatSpatial, a schema enforced orchestration framework in which an LLM selects versioned tools and parameters rather than executable code. Built on the Model Context Protocol, ChatSpatial integrates 65 methods across 15 analytical categories into 20 typed tools with curated defaults. In two cancer case studies, ChatSpatial coordinated cross ecosystem workflows that recovered the analytical structure and central conclusions of published ovarian cancer and oral squamous cell carcinoma analyses. Across three LLMs, schema enforcement raised parameter validity from 20% to 98% and stabilized method selection. In end to end benchmarks against two code generation alternatives, it yielded higher execution success and output concordance.

## 1 Introduction

As spatial transcriptomics platforms have matured, the analytical bottleneck has shifted from data generation to workflow composition [1, 2]. A single study may require spatial domain identification, deconvolution, cell–cell communication analysis, and pathway enrichment—each step drawing on specialized methods distributed across partially incompatible software ecosystems [3], principally Python (Scanpy/AnnData [4]) and R (Seurat [5]). The result is that a biologically coherent analysis often becomes an exercise in environment management, data conversion, and manual stitching of intermediate outputs.

The first-principles problem is therefore not simply that spatial transcriptomics lacks methods. Rather, analytical intent must be translated into a sequence of executable, inspectable operations across methods, languages, and data structures. Existing strategies address parts of this translation problem. Bridge packages such as `zellkonverter` handle format conversion; interactive co-pilots assist with individual analysis steps [6–8]; and LLM-based spatial agents use tool libraries or generated code to automate broader workflows [9, 10]. These approaches typically trade off among low-friction interaction, cross-ecosystem execution, and workflow-level transparency. Unconstrained code generation is flexible, but it expands the output space to package APIs, syntax, imports, and undocumented intermediate choices; direct programming is transparent, but it places the integration burden on the researcher.

Here, we present ChatSpatial, a schema-enforced orchestration framework for spatial transcriptomics. The central design choice is to separate flexible intent interpretation from deterministic scientific execution: the LLM maps a user request to versioned tool schemas and parameter fields, while established Python and R methods perform the computation through validated wrappers. Built on the Model Context Protocol (MCP) [11], this design constrains each request to a defined parameter space and records the resulting workflow as structured tool calls, parameter values, data identifiers, and software versions. Method suitability, defaults, and data-format requirements are encoded in schema descriptions, giving the model visible guidance without allowing it to invent executable code.

We evaluate ChatSpatial at three levels. Published-data case studies test whether the framework preserves analytical structure and central biological interpretation in realistic workflows; controlled experiments test whether schema enforcement stabilizes tool selection, parameter specification, and execution outcomes; and functional edge cases test whether ambiguity and failures are surfaced rather than silently hidden.

## 2 Results

### 2.1 Schema-constrained orchestration architecture

The current implementation exposes 65 methods across 15 analytical categories through 20 MCP tools (Figure 1a–b). Each tool accepts typed parameter fields validated by Pydantic models before any analysis code runs. Of the 441 total parameters, 358 (81.2%) are constrained through Literal enumerations, numeric bounds, or default values; the remaining 83 (18.8%) accept free-text input, primarily dataset identifiers and column names that reference loaded data structures (Methods §4.1). Each invocation is recorded as a structured tuple of tool name, parameter values, data identifiers, and software versions.

**Fig. 1.**
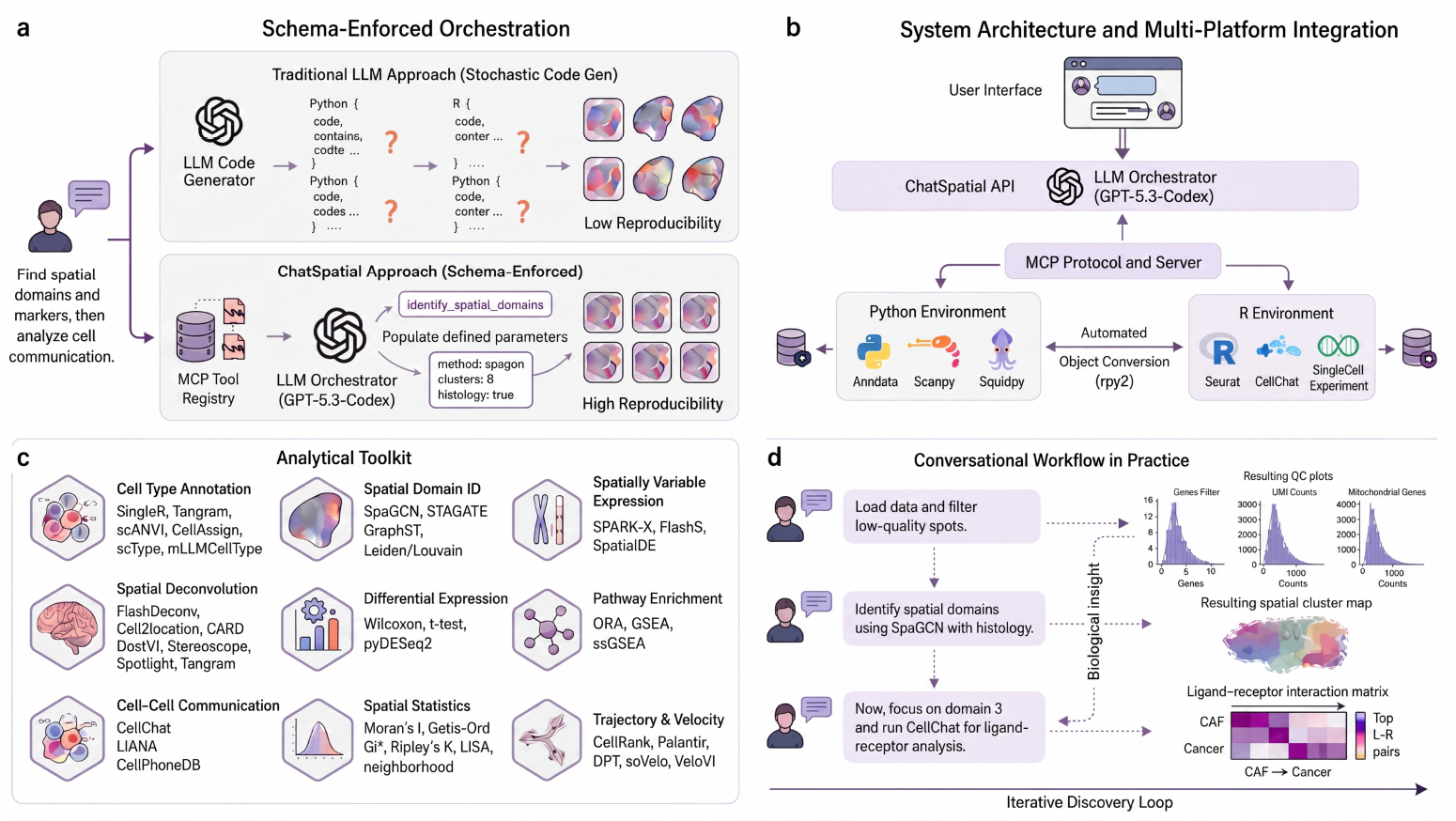
Overview of the ChatSpatial schema-enforced orchestration architecture. (**a**) Unconstrained code generation versus schema-enforced orchestration. Top: free-form LLM output must specify package APIs, syntax, imports, and execution details, creating a broad and variable output space. Bottom: ChatSpatial constrains model output to versioned MCP tool schemas, limiting the model-dependent step to tool and parameter selection. (**b**) System architecture and cross-ecosystem integration. User queries pass through the ChatSpatial API to an LLM orchestrator, which invokes tools via the MCP protocol and server layer. Automated object conversion (rpy2) bridges the Python environment (AnnData, Scanpy, Squidpy) and R environment (Seurat, CellChat, SingleCellExperiment), supporting cross-ecosystem workflows in the tested analyses. (**c**) Representative analytical categories from the implemented toolkit, including cell type annotation, spatial domain identification, spatially variable expression, spatial deconvolution, differential expression, pathway enrichment, cell–cell communication, spatial statistics, and trajectory or velocity analysis. The full method list is provided in Supplementary Table 2. (**d**) Conversational workflow in practice. Three sequential prompts—data loading with quality control, spatial domain identification, and ligand–receptor analysis—illustrate how structured analysis outputs are produced through iterative natural-language interaction.

### 2.2 Cross-ecosystem implementation breadth

The best-benchmarked methods for different analytical steps often live in different software ecosystems. ChatSpatial bridges Python workflows built around AnnData, Scanpy, and Squidpy with R methods such as Seurat, CellChat, RCTD, and SingleCellExperiment through wrapper-level conversion via `rpy2`, allowing a single recorded workflow to combine methods that would otherwise require manual data export, format conversion, and environment management.

The resulting toolkit spans 15 analytical categories—from spatial domain identification and deconvolution to cell communication, trajectory analysis, and copy-number inference (Figure 1c; Table 1; Supplementary Table 2)—following established computational taxonomies [3, 12]. Within each category, methods were selected to provide benchmarked or widely used representatives [13–19]; multiple alternatives are included so that users can compare approaches with distinct assumptions.

**Table 1.** Platform compatibility matrix: implemented analytical module support across representative spatial transcriptomics technologies. **Legend:** ✓✓= commonly used or wrapper-recommended for this data type, ✓= implemented support, ◦ = optional or assay-dependent, *−* = not required/not applicable. Cell-resolution technologies (MERFISH, Xenium, seqFISH, CosMx) do not require deconvolution as they provide native single-cell data. Spot-based platforms (Visium, Slide-seq) benefit from deconvolution to infer cell-type compositions. RNA velocity requires spliced/unspliced counts or equivalent assay fields. Support denotes implemented wrapper compatibility; biological validation was performed on representative workflows rather than every method–platform combination.

| Module | Visium | Visium<br>HD | MERFISH<br>Xenium | seqFISH<br>CosMx | Slide<br>-seq | STAR<br>map |
| --- | --- | --- | --- | --- | --- | --- |
| **Preprocessing** | ✓ | ✓ | ✓ | ✓ | ✓ | ✓ |
| **Multi-Sample**<br>**Integration** | ✓ | ✓ | ✓ | ✓ | ✓ | ✓ |
| **Cell Type Anno-**<br>**tation** | ✓ | ✓ | ✓ | ✓ | ✓ | ✓ |
| **Differential**<br>**Expression** | ✓ | ✓ | ✓ | ✓ | ✓ | ✓ |
| **Spatial Domain**<br>**ID** | ✓✓ | ✓ | ✓ | ✓ | ✓ | ✓ |
| **Spatial Variable**<br>**Genes** | ✓ | ✓ | ✓ | ✓ | ✓ | ✓ |
| **Cell Communi-**<br>**cation** | ✓ | ✓ | ✓✓ | ✓✓ | ✓ | ✓ |
| **Deconvolution** | ✓✓ | ○ | – | – | ○ | ○ |
| **CNV Analysis** | ✓ | ✓ | ✓ | ✓ | ✓ | ✓ |
| **Trajectory Anal-**<br>**ysis** | ✓ | ✓ | ✓✓ | ✓✓ | ✓ | ✓ |
| **RNA Velocity** | ✓ | ✓ | ✓ | ✓ | ✓ | ✓ |
| **Enrichment**<br>**Analysis** | ✓ | ✓ | ✓ | ✓ | ✓ | ✓ |

We tested implementation coverage across 31 scenarios spanning datasets from low-density STARmap (300 spots, 150 genes) to high-resolution Xenium (150K cells), as well as multi-sample Visium workflows requiring integration (§2.5, Supplementary Table 3). The controlled reproducibility analysis is presented separately below.

### 2.3 Workflow fidelity on published spatial datasets

We next asked whether the framework can preserve the analytical structure of realistic spatial transcriptomics studies. The evaluation criterion is workflow fidelity rather than exact pipeline replication: a successful reproduction should coordinate the same class of analytical steps, expose intermediate choices and outputs, and recover the central biological interpretation, even when wrapper-level implementations differ from the original publication. We selected two published cancer studies that stress different parts of the architecture: OSCC for spatial-domain, deconvolution, and R-based cell communication chaining; and HGSOC for multi-sample deconvolution, copy-number inference, and cross-patient summarization.

#### 2.3.1 Oral Cancer: Tumor Architecture and Cell-Cell Communication

We used ChatSpatial to reproduce the main analytical workflow of Arora et al. [21], who characterized the distinct transcriptional architectures of the Tumor Core (TC) and Leading Edge (LE) in oral squamous cell carcinoma (OSCC) through a multi-step pipeline involving cohort integration, unsupervised spatial domain identification, reference-based deconvolution, and ligand-receptor analysis.

##### Step 1: Unsupervised Spatial Domain Discovery and Cohort Integration

The workflow began with identifying the primary spatial domains across all 12 patient samples: *“Load all 12 OSCC samples, filter low-quality spots (<200 genes), and identify spatial domains using Leiden clustering with spatially-aware neighbor graphs*.*”* This single instruction coordinated cohort ingestion and unsupervised spatial domain identification. Although the original study used Louvain clustering with Seurat’s CCA-based integration on malignant spots only, the ChatSpatial workflow recovered a consistent domain structure: a Tumor Core (TC) enriched for keratinization and epithelial differentiation markers (e.g., *SPRR1B, CLDN4, SPRR2E*), a Transitory state marked by *KRT6C*, and a Leading Edge (LE) enriched for partial-EMT and invasion markers (e.g., *FN1, LAMC2, ITGA5*).

##### Step 2: Deconvolution Reveals Conserved Microenvironment Composition

To assess cellular composition, deconvolution was performed using the HNSCC single-cell reference from Puram et al. [20]: *“Deconvolve cell types in the identified TC and LE domains using CARD, utilizing the HNSCC single-cell reference from Puram et al*.*”* ChatSpatial processed the reference and executed CARD across the spatial data. The resulting composition estimates recapitulated the dominant immune-stromal pattern reported by Arora et al.: the Leading Edge was enriched for macrophages (11/12 samples, 92%; 95% CI 62–100%) and fibroblasts/CAFs (10/12, 83%; 95% CI 52–98%) relative to the Tumor Core, while cancer cells showed TC-directional enrichment in 7/12 samples (58%; 95% CI 28–85%), below the 75% consensus threshold (Figure 2a–e).

**Fig. 2.**
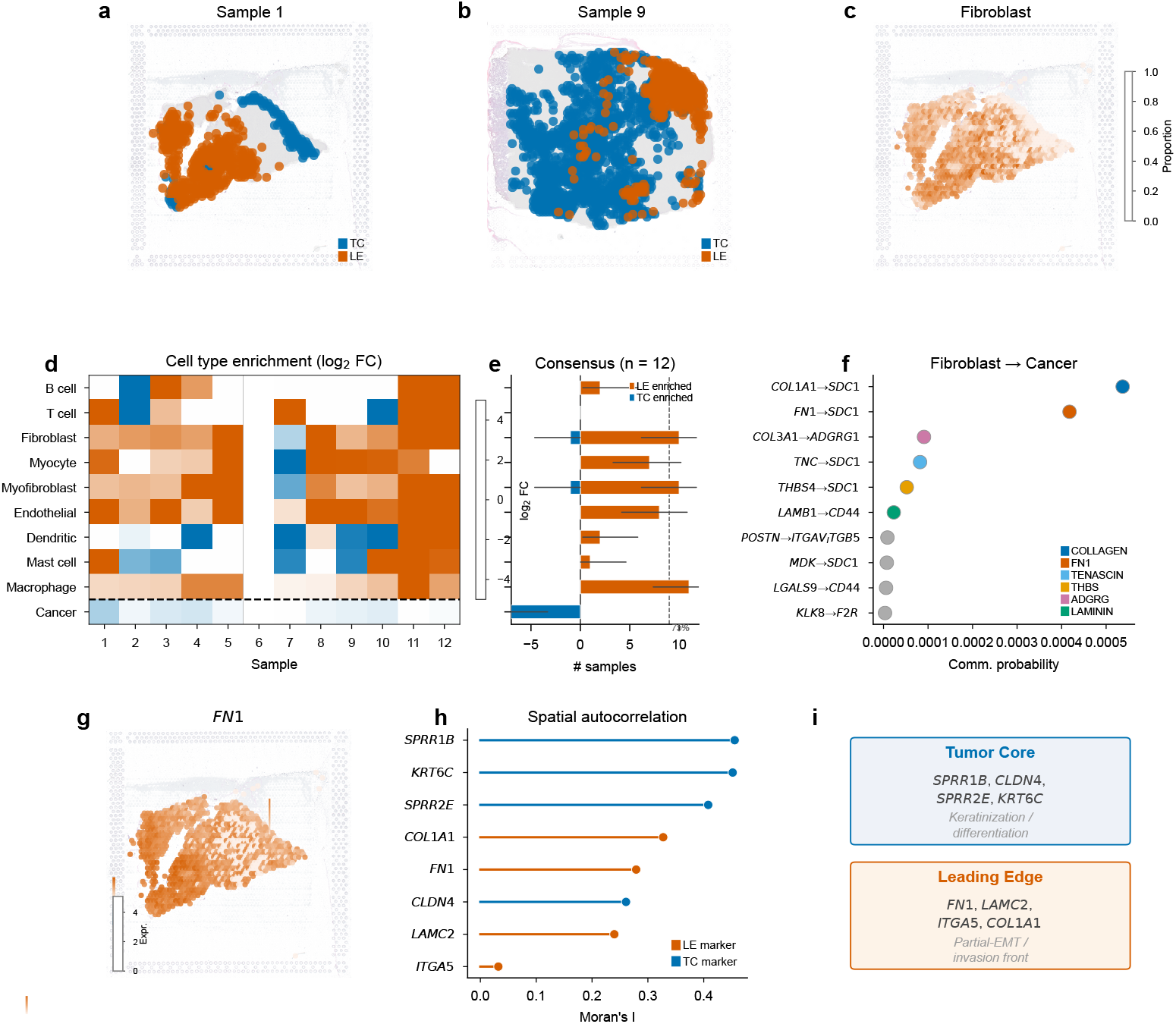
ChatSpatial reproduces the analytical structure of a published OSCC tumor architecture workflow through conversational orchestration. (**a, b**) H&E tissue sections of two representative OSCC samples (Sample 1 and Sample 9 from GSE208253) with spatial domain annotations: Tumor Core (TC, blue) and Leading Edge (LE, orange). Spatial domains were identified by Leiden clustering on spatially-aware neighbor graphs and classified using marker gene scoring. (**c**) Spatial deconvolution map (Sample 1) showing fibroblast/CAF proportion at each Visium spot (CARD with HNSCC scRNA-seq reference [20]), revealing fibroblast concentration at the leading edge. (**d**) Per-sample cell type enrichment between TC and LE across 12 samples (11 evaluable; Sample 6 excluded for lacking LE spots). Color indicates within-sample log_2_ fold change (LE/TC) of mean deconvolution proportion. Sample-level inference (Wilcoxon signed-rank on per-patient log_2_FC; *n* = 11, treating each patient as the biological replicate) is reported in Supplementary Table 1. (**e**) Consensus enrichment summary (*n* = 12 patients) with exact binomial (Clopper–Pearson) 95% CIs (horizontal lines). A sample is counted as enriched if its within-sample test is significant (BH-FDR *q <* 0.05) and | log_2_FC | *>* 0.25. Dashed line: 75% threshold (≥9/12 samples). Macrophages (11/12, 92%) and fibroblasts (10/12, 83%) show robust LE enrichment; cancer cells show TC-directional enrichment in 7/12 samples (58%). (**f**) CellChat ligand-receptor analysis identifying top Fibroblast → Cancer interactions colored by signaling pathway. Directional ECM-syndecan interactions (COL1A1 → SDC1, FN1 → SDC1) align with the ECM-associated tumor–stroma axis proposed in the original study [21]. (**g**) Spatial expression of *FN1* (Sample 1), confirming localization of the CellChat-identified ligand to the LE domain. (**h**) Moran’s *I* spatial autocorrelation for key TC and LE marker genes in Sample 1. TC marker *SPRR1B* and LE markers *COL1A1* and *FN1* ranked within the top 0.2% of spatially patterned genes in the tested gene set. (**i**) Summary of TC and LE marker gene signatures. TC markers are associated with keratinization and epithelial differentiation; LE markers are associated with partial-EMT and invasion front programs.

Sample-level Wilcoxon signed-rank tests (*n* = 11 patients, treating each patient as the biological replicate; Sample 6 excluded for lacking LE spots) confirmed the robustness of these patterns: significant LE enrichment for macrophages (*p*_FDR_ = 0.003), fibroblasts (*p*_FDR_ = 0.005), CAFs (*p*_FDR_ = 0.027), and endothelial cells (*p*_FDR_ = 0.003), and significant TC enrichment of cancer cells (*p*_FDR_ = 0.003; Supplementary Table 1). Endothelial cells (8/12, 67%) and myocytes (7/12, 58%) showed consistent but sub-threshold LE-directional enrichment. The strongest signals—macrophage and fibroblast LE enrichment—align with the fibrovascular niche described in the original study; weaker signals may reflect biological heterogeneity across patients.

##### Step 3: Cross-Ecosystem Ligand–Receptor Analysis

The final step examined candidate tumor–stroma interactions by invoking native R tools through ChatSpatial’s cross-ecosystem bridge. Following the original study’s focus on the invasive front, the prompt was: *“Focus on the Leading Edge (LE) domain and perform cell communication analysis between CAFs and cancer cells using CellChat. Specifically, look for interactions involving ECM ligands*.*”*

Through cross-ecosystem invocation of the R-based CellChat package, ChatSpatial identified 142 significant Fibroblast → Cancer interactions spanning six signaling pathways (COLLAGEN, FN1, TENASCIN, THBS, LAMININ, and ADGRG). The top-ranked interactions—COL1A1 → SDC1, COL1A2 → SDC1, and FN1 → SDC1—aligned with the ECM–syndecan tumor–stroma interaction axis proposed in the original study (Figure 2f). Spatial expression mapping of *FN1* confirmed its localization to the LE domain (Figure 2g).

#### Exploratory Cross-Method Spatial Autocorrelation

As a follow-up analysis not reported by Arora et al., spatial autocorrelation (Moran’s *I*) was used to ask whether the CellChat-prioritized ECM ligands also showed strong spatial patterning. A single prompt—*”Run spatial autocorrelation on this sample and check whether FN1 and COL1A1 are spatially variable”*—triggered ChatSpatial to compute Moran’s *I* in Sample 1 (squidpy; Benjamini–Hochberg FDR correction). Both LE markers ranked among the most spatially patterned genes in the tested set: *FN1* (rank 12/15,215, *I* = 0.51) and *COL1A1* (rank 18, *I* = 0.49). TC markers showed similarly strong spatial patterning, including *SPRR1B* (rank 8, *I* = 0.53), *KRT6C* (rank 37, *I* = 0.46), and *SPRR2E* (rank 41, *I* = 0.46). These patterns were consistent across the cohort: *FN1* spatial autocorrelation remained strong across 11 samples (mean *I* = 0.28, 95% CI [0.18, 0.38]), as did *COL1A1* (*I* = 0.33 [0.24, 0.42], *n* = 11) and *SPRR1B* (*I* = 0.46 [0.35, 0.56], *n* = 9). This cross-method connection—from cell communication signatures to spatial gene expression rankings—required a single follow-up prompt (Figure 2g–h).

#### 2.3.2 Ovarian Cancer: Multi-Sample Subclone and Microenvironment Analysis

Denisenko et al. profiled eight HGSOC patient samples spanning different chemotherapy response categories, using an intricate computational pipeline to identify tumor subclones and characterize their microenvironments [22]. ChatSpatial was applied to all eight samples (P1–P8) to recover the main analytical structure of this workflow (Figure 3).

**Fig. 3.**
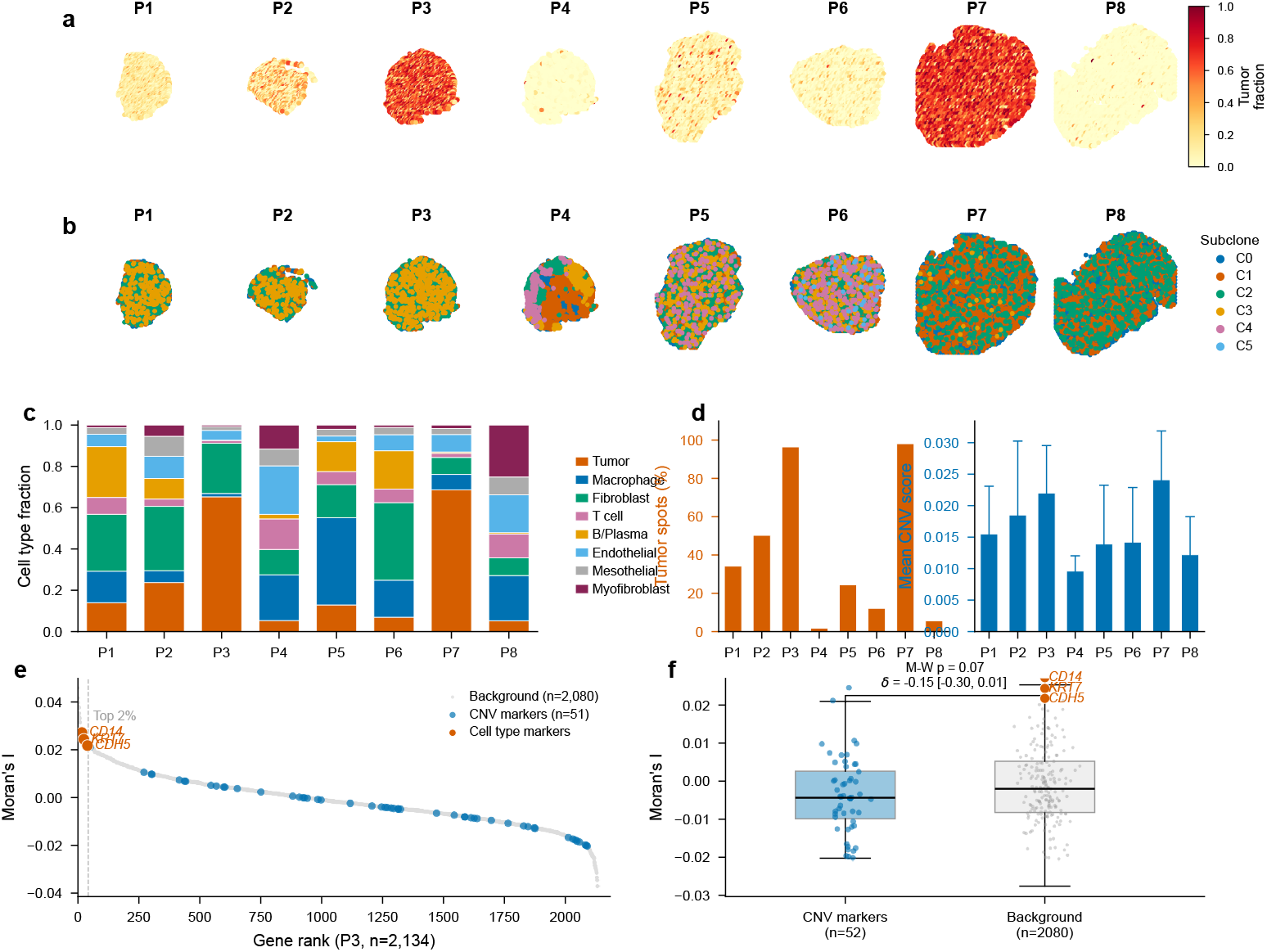
ChatSpatial recovers the analytical structure of a published HGSOC subclone analysis through a multi-sample conversational workflow. Reanalysis of Denisenko et al. [22] using publicly available HGSOC Visium data (GSE211956, 8 patient samples: P1–P8). (**a**) Tumor cell fraction spatial maps from RCTD deconvolution across all patients. The publicly deposited scRNA-seq reference contains 8 consolidated cell types (the original study used a 12-type annotation with 5 fibroblast subtypes; see Methods). Color intensity indicates tumor fraction (yellow = low, red = high). Tumor-enriched spots (RCTD tumor weight ≥ 0.15) were identified for downstream analysis. (**b**) infercnvpy-based copy number cluster mapping reveals spatially distinct CNV profiles across patients. Multiple CNV clusters (3–6 per patient) were identified in all 8 samples via Leiden clustering on the inferred CNV matrix, consistent with the original study’s observation of genomic heterogeneity validated by whole-genome sequencing. Note: the original study used manual dendrogram-based splitting; automated Leiden clustering may yield different partition granularities. (**c**) Cell type composition across patients from RCTD deconvolution, showing variable proportions of tumor cells, macrophages, fibroblasts, T cells, B/plasma cells, endothelial cells, mesothelial cells, and myofibroblasts. (**d**) Quantitative summary of tumor burden (vermillion bars, left axis: percentage of tumor-enriched spots) and genomic instability (blue bars, right axis: mean CNV score *±* SD across spots) across patients, enabling cross-patient comparison of microenvironment dynamics. (**e**) Spatially variable gene ranking for patient P3 by Moran’s *I*, with cell type marker genes (*CD14, KRT7, CDH5* ; red) and CNV-cluster marker genes (blue) highlighted. Cell type markers rank among the strongest spatially patterned genes in this sample, while CNV markers are distributed more broadly across the ranking. (**f**) Moran’s *I* distributions for CNV-cluster marker genes versus background genes (Mann–Whitney *p* = 0.07; Cliff’s *δ* = *−*0.15 [95% CI *−*0.30, 0.01]). The difference is not statistically significant; this panel illustrates the cross-method chaining capability rather than a claimed biological effect.

##### Step 1: Multi-Sample Integration and Tumor Identification

The workflow began with processing all eight patient samples: *“Load the HGSOC Visium samples from GSE211956 and deconvolve cell types using RCTD with the study’s scRNA-seq reference*.*”* This single instruction triggered batch processing across the cohort, applying RCTD deconvolution [23] with the study’s scRNA-seq reference (8 cell types in the publicly deposited version; the original study used a finer 12-type annotation that subdivided fibroblasts into 5 subtypes). ChatSpatial automatically identified tumor-enriched spots (RCTD tumor weight ≥ 0.15, following the original study’s convention) across all samples, enabling systematic downstream analysis (Figure 3a,c).

##### Step 2: Subclone Discovery via Copy Number Inference

Candidate subclonal populations were identified through copy number inference: *“For each sample, run infercnvpy using low-tumor spots as the normal reference to identify regions with distinct copy number alteration profiles*.*”* ChatSpatial executed infercnvpy [24, 25] per sample, using spots with low RCTD tumor weights (*<* 0.15) as the reference profile—a critical methodological detail from the original study. Leiden clustering on the inferred CNV matrix revealed multiple spatially distinct CNV clusters in all 8 patients (3–6 clusters per sample), consistent with the original study’s finding of pervasive copy number heterogeneity across the HGSOC cohort (Figure 3b).

##### Step 3: Cross-Patient Microenvironment Comparison

Cross-patient microenvironment comparison was the final analytical step: *“Summarize the cell type composition and tumor burden across all patients. Which patients show the highest genomic instability?”* The resulting cross-patient analysis revealed substantial heterogeneity in both tumor burden (ranging from 2% to 98% tumor-enriched spots) and genomic instability (mean CNV scores 0.010–0.024, SD 0.002–0.012 across spots within each patient; Figure 3d), with patients harboring the highest tumor fractions (P3, P7) also exhibiting complex subclonal architectures in this cohort (Figure 3c–d). These patterns are consistent with the original study’s observation that intratumor heterogeneity varies substantially across patients and may influence differential treatment response.

##### Exploratory Cross-Method Spatial Analysis

Patient P3—a tumor-rich sample with multiple CNV clusters—was selected for a cross-method follow-up: a single conversational prompt performed spatial autocorrelation (Moran’s *I* [26]) across expressed genes. Cell type markers from the RCTD deconvolution, including *CD14* (monocyte/macrophage), *KRT7* (epithelial/tumor), and *CDH5* (vascular endothelial), ranked among the strongest spatially patterned genes in this sample. CNV-cluster marker genes were more dispersed across the spatial-autocorrelation distribution, though the difference was not statistically significant (Mann–Whitney *p* = 0.07; Cliff’s *δ* = *−* 0.15 [95% CI *−* 0.30, 0.01]; Figure 3e–f). This analysis required linking deconvolution, CNV inference, and spatial statistics in a single conversational session.

Together, these case studies show that typed tool schemas can preserve the analytical structure of published workflows while enabling low-friction cross-method exploration.

### 2.4 Controlled reproducibility experiments

A separate question is whether schema enforcement *itself* —rather than any structured tool-calling interface—causally contributes to consistency. To isolate this effect, we compared three conditions: *full schema* (typed parameters with Literal enumerations, bounds, defaults, and natural-language descriptions—the current ChatSpatial design), *bare schema* (types and enumerations retained, all descriptions removed), and *no schema* (tool names with one-sentence capability summaries, no typed parameters). All three conditions required the same JSON response format, so any performance difference reflects schema content, not output structure. We evaluate two levels of reproducibility: *invocation reproducibility* —the probability that identical analytical intent produces identical tool calls and schema-valid parameters across independent LLM samples—and *output reproducibility* —the concordance of analytical results (Adjusted Rand Index, Jaccard similarity, Pearson correlation) across such invocations.

#### Invocation-level ablation

Eight representative prompts were presented to three LLMs under each condition (720 total trials; see Methods §4.5 for model selection rationale and experimental design). Each response was validated against ChatSpatial’s Pydantic parameter models—the same validation that gates real tool execution.

Full-schema invocations passed Pydantic validation in 98.3% of trials (Figure 4a). Bare schema achieved 93.8%, indicating that type constraints alone capture most of the validation benefit. Without any schema, the rate fell to 20.4%, consistent across all three models (Claude Haiku 4.5: full 100%, no-schema 23.8%; GPT-5.4: 100% to 16.2%; Gemini 2.5 Flash: 95.0% to 21.2%). Tool selection remained 100% consistent under all conditions; the constraint gap lies entirely in parameter specification.

**Fig. 4.**
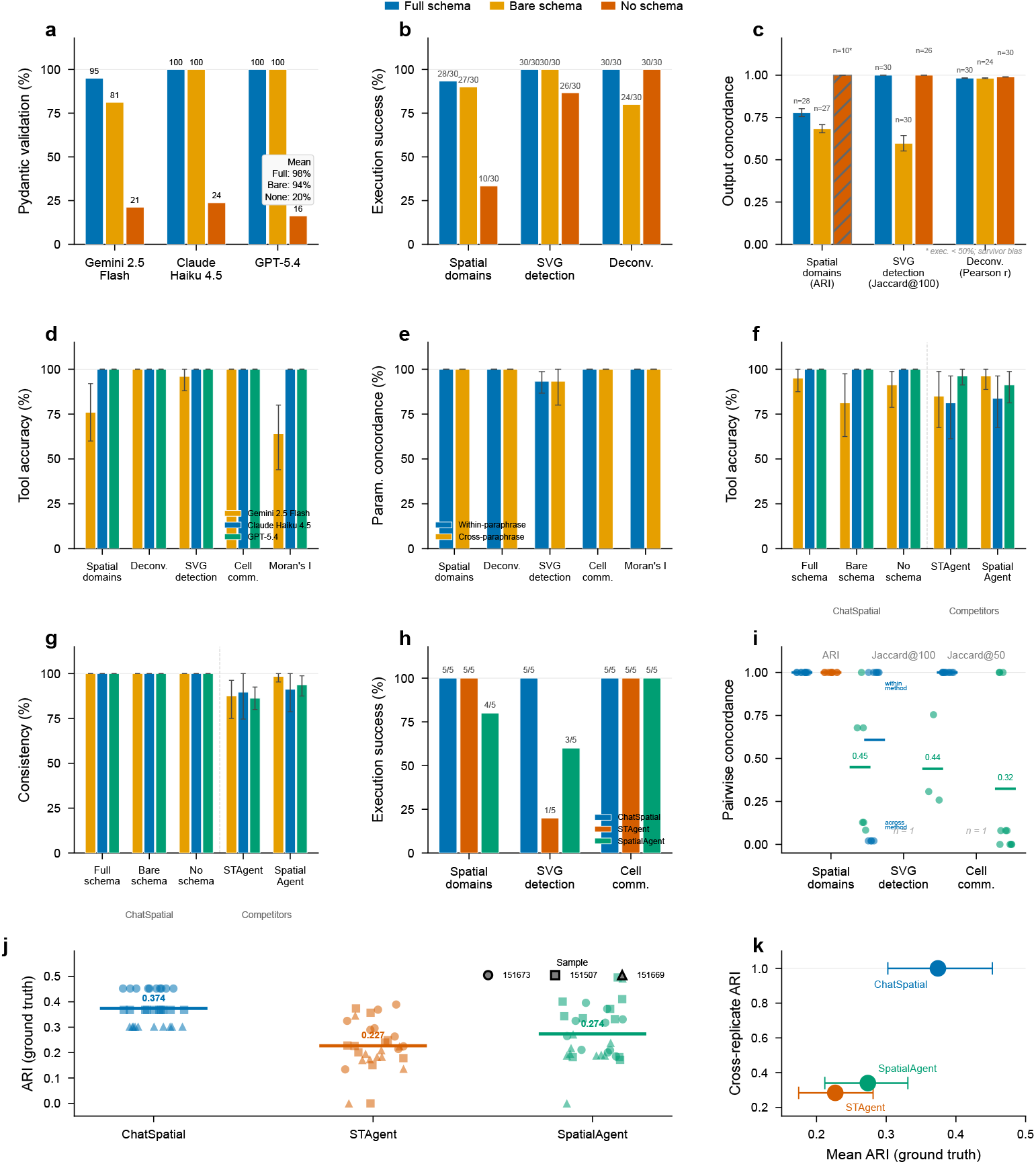
Schema-enforcement ablation, prompt sensitivity, and cross-system comparison. *Top row* (ablation; 720 invocation-level trials, 270 end-to-end trials): (**a**) Pydantic validation rate under full schema (types + enumerations + descriptions), bare schema (types + enumerations only), and no schema (tool names only). Each bar represents the validation rate across 80 trials (8 prompts × 10 replicates). (**b**) End-to-end execution success for three analytical tasks on an OSCC Visium dataset. Labels show successful/total trials aggregated across models. (**c**) Cross-model output concordance for successful trials, measured by Adjusted Rand Index (spatial domains), Jaccard@100 (SVG detection), and Pearson *r* (deconvolution). Error bars: bootstrap 95% CIs (10,000 resamples). Hatched bars indicate conditions with *<*50% execution success, where concordance is inflated by survivor bias. *Middle row* (prompt sensitivity and cross-system tool selection): (**d**) Tool selection accuracy across semantically equivalent paraphrases per prompt group and model (375 trials: 5 prompt groups × 5 paraphrases × 3 LLMs × 5 replicates, *T* = 1.0, full-schema condition). (**e**) Constrained parameter concordance: within-paraphrase consistency (single prompt, multiple replicates) versus cross-paraphrase consistency (five paraphrases). (**f**) Cross-system tool selection accuracy (1,200 total trials across five conditions). Dashed line separates ChatSpatial conditions from competitor system contexts. *Third row* (cross-system consistency and end-to-end benchmark): (**g**) Cross-replicate tool consistency across systems. (**h**) End-to-end execution success comparing ChatSpatial, STAgent, and SpatialAgent on a shared Visium dataset (human lymph node, Claude Sonnet 4, 5 replicates per system per task). Pairwise output concordance among successful replicates: ARI for spatial domains, Jaccard@100 for SVG rankings, Jaccard@50 for ligand-receptor pair overlap. *Bottom row* (ground-truth benchmark): (**j**) ARI against expert-annotated cortical layers on DLPFC Visium data (3 samples *×* 3 systems *×*10 replicates). Points show individual trials; bars show system means. (**k**) Accuracy– consistency tradeoff: mean ground-truth ARI versus mean cross-replicate ARI for each system.

#### End-to-end execution

We next tested whether this validation gap has functional consequences by extending the ablation to three executable tasks—spatial domain identification (Leiden clustering), spatially variable gene detection (FlashSV), and cell-type deconvolution (FlashDeconv)—on an oral squamous cell carcinoma Visium dataset (270 total trials; Figure 4b). Each trial produced an LLM invocation, validated parameters via Pydantic, and executed the tool on a fresh data copy if valid.

Execution success depended on condition and task complexity. Spatial domain identification, which requires specifying method, resolution, and clustering parameters, succeeded in 93% of full-schema attempts (28/30) but only 33% without schema (10/30). Spatially variable gene detection achieved 100%, 100%, and 87% across conditions. Deconvolution—where most parameters have robust defaults—achieved 100%, 80%, and 100%; consistent with this, the output concordance analysis below shows that deconvolution results were nearly identical across all three conditions (Pearson *r* ≥ 0.98), suggesting that the algorithm’s parameter space is narrow enough for unguided LLMs to find valid configurations.

#### Output concordance

Among successful trials, we measured pairwise cross-model concordance: Adjusted Rand Index (ARI) for cluster labels, Jaccard similarity of top-100 ranked genes, and Pearson correlation of cell-type proportions (Figure 4c). Full-schema SVG detection yielded complete cross-model agreement (Jaccard@100 = 1.0 [95% CI: 1.0–1.0]); bare-schema concordance dropped to 0.60 [0.55–0.64], reflecting method-choice divergence when descriptions were removed. Deconvolution concordance was high across all conditions (Pearson *r >* 0.98), consistent with the limited parameter sensitivity of the underlying algorithm. For spatial domains, the 10 no-schema trials that passed validation achieved ARI = 1.0, but this reflects survivor bias: only invocations with valid default parameters could execute, yielding mechanically identical outputs from a population where 67% failed entirely. The same survivor effect applies to no-schema SVG detection (Jaccard@100 = 1.0 from 26/30 trials that passed): surviving invocations converged on the same method and parameters, inflating apparent concordance.

In sum, typed constraints raised parameter validity from 20% to 98% (omnibus Kruskal–Wallis *p <* 0.001 for all three tasks). The execution gap was largest for parameter-sensitive tasks (spatial domains: 93% vs. 33%) and smallest where algorithms have robust defaults (deconvolution: Pearson *r* ≥ 0.98 across all conditions; full vs. bare *p* = 0.22, n.s.). Bare schema recovered ∼ 94% validation, but cross-model method-choice concordance required descriptive guidance (SVG Jaccard: full 1.0 vs. bare 0.60, Mann–Whitney *p <* 0.001). Full pairwise contrasts are in the reproducibility repository.

#### Case-study workflow concordance

The ablation above tests generic prompts; a separate question is whether this consistency extends to the specific workflows in §2.3. We repeated the OSCC CARD deconvolution (Step 2) ten times per model under full-schema and no-schema conditions, using the three ablation models plus Claude Sonnet 4.5—the model used for the case studies—on two patient samples (Samples 1 and 9). CARD deconvolution is deterministic given fixed parameters, so any output variation reflects parameter-selection variation induced by the schema condition. Under full schema, all four models produced valid invocations in every trial (80/80), each selecting the same CARD parameters and yielding identical proportion matrices (pairwise Pearson *r* = 1.000 [95% CI: 1.000–1.000]; 780 cross-model pairs per sample). Under no schema, every model correctly identified the tool and supplied otherwise valid arguments, yet all 80 trials failed Pydantic validation because the models used the official method name `CARD` rather than the lowercase enumeration value `card`. These results confirm that the deconvolution findings in Figure 2 are reproducible across models and are not contingent on a single LLM invocation.

#### Prompt sensitivity

The ablation above varies the schema while holding the user prompt fixed. A complementary question is whether semantically equivalent rephrasings of the same request produce consistent tool selection and parameters. To test this, we generated five paraphrases for each of five representative tasks (spatial domains, deconvolution, SVG detection, cell communication, Moran’s I), presented each to all three LLMs under the full-schema condition with five replicates at *T* = 1.0 (375 total trials). Tool selection accuracy across paraphrases was 95.7%. Claude Haiku 4.5 and GPT-5.4 each achieved 100%; Gemini 2.5 Flash reached 87.2%, with all failures due to response-format parsing rather than incorrect tool choice. Cross-paraphrase constrained-parameter concordance was 98.7%, comparable to the 98.0% within-paraphrase consistency observed in the ablation (Figure 4d,e). These results indicate that under schema enforcement, tool selection and constrained parameter values are robust to natural-language variation in user intent expression.

#### Cross-system invocation comparison

We next asked whether this advantage extends beyond ChatSpatial’s own catalog. We extracted the system context—tool catalog, system prompt, and routing instructions— from STAgent [9] (8 tools, including a general-purpose code-execution environment) and SpatialAgent [10] (72 specialized tools across seven analytical categories), and presented the same eight prompts to the same three LLMs under each system’s context at *T* = 1.0 (480 new trials; 1,200 total across five conditions; Figure 4f,g). ChatSpatial’s full-schema condition achieved 98.3% tool selection accuracy [95% CI: 95.4–100.0] with 100% cross-replicate consistency, compared with 87.5% [77.9–95.4] and 87.8% consistency for STAgent, and 90.4% [83.3–96.7] and 94.5% consistency for SpatialAgent. The accuracy gap was task-dependent: STAgent routes most analytical queries through a general-purpose code-execution tool, leaving tool choice architecturally underspecified for domain-specific tasks; SpatialAgent’s larger catalog provides more targeted options but introduces selection ambiguity where no exact match exists (e.g., CNV inference: 60% accuracy in both systems). This comparison isolates single-step tool selection; code generation quality and multi-turn execution are outside its scope.

#### End-to-end execution benchmark

We ran all three systems on a shared Visium dataset (human lymph node, 10x Genomics; not used in any case study) with identical prompts and the same LLM (Claude Sonnet 4, *T* = 1.0, 5 replicates per system per task; Figure 4h,i). Claude Sonnet 4 was chosen for cross-system compatibility (the case-study analyses used Claude Sonnet 4.5; see Methods). ChatSpatial completed all 15 trials; STAgent completed 11/15 (73%), with all 4 failures in SVG detection due to recursion-limit exhaustion; SpatialAgent completed 12/15 (80%), with failures from recursion limits and timeouts.

Output concordance separated the systems further. For spatial domains, ChatSpatial and STAgent both achieved ARI = 1.0 across all successful replicate pairs; SpatialAgent produced more variable clusterings (mean ARI = 0.45 [0.20–0.74]), reflecting code-generation variation in preprocessing and resolution parameters. For SVG detection, ChatSpatial replicates selecting the same method showed Jaccard@100 = 1.0, though at *T* = 1.0 the LLM chose a different detection method in one replicate, producing a bimodal concordance distribution; STAgent’s single successful replicate precluded pairwise comparison. For cell–cell communication, ChatSpatial achieved Jaccard@50 = 1.0 across all 10 replicate pairs; SpatialAgent produced variable ligand– receptor pair lists (mean Jaccard@50 = 0.32 [0.05–0.62]); STAgent completed all trials but its interactive confirmation requirement prevented automated output extraction.

The SVG result illustrates a residual source of variation even within the constrained architecture: when the LLM selected different detection methods across replicates, pairwise concordance dropped accordingly.

#### Ground-truth benchmark

The end-to-end benchmark above measures execution success and cross-replicate concordance; a complementary question is whether the outputs are biologically correct. We evaluated all three systems on the DLPFC benchmark [27]—a standard evaluation dataset for spatial domain methods where expert-annotated cortical layers (L1–L6 and white matter) serve as ground truth. Each system received an open-ended prompt (“Identify spatial domains in this dataset”) without specifying the method, resolution, or number of domains, and ran 10 replicates on three samples from distinct donors (90 total trials; Claude Sonnet 4, *T* = 1.0). Ground-truth concordance was measured by ARI against expert annotations.

ChatSpatial achieved the highest accuracy (mean ARI = 0.374 [0.302–0.452]) with perfect cross-replicate consistency (pairwise ARI = 1.000; 135 pairs), selecting SpaGCN with *n* = 7 domains deterministically in all 30 trials—producing identical outputs across replicates within each sample. SpatialAgent achieved ARI = 0.274 [0.211– 0.331] with cross-replicate ARI = 0.340; STAgent achieved ARI = 0.227 [0.174–0.281] with cross-replicate ARI = 0.284. (All 95% CIs: hierarchical bootstrap over three samples.) All three systems completed every trial (90/90). The accuracy advantage reflects both method choice—the schema-guided selection of SpaGCN with histology-aware spatial modeling—and the deterministic execution that eliminates run-to-run variation. Code-generation systems produced correct but variable clusterings, with different preprocessing pipelines and resolution parameters across replicates lowering both accuracy and consistency.

### 2.5 Failure modes and human-in-the-loop refinement

Finally, we evaluated how the framework behaves under functional edge cases, ambiguous user intent, and observed failure conditions. We tested 31 predefined scenarios covering data handling and preprocessing (5), core spatial analysis (11), conversational workflows (5), scalability stress tests (7), and known limitations (3) (Supplementary Table 3). Of these, 25 produced a valid analysis result or an explicit diagnostic on the first turn; three involving underspecified prompts required one round of conversational clarification before succeeding. The remaining three scenarios exposed genuine limitations: an unsupported method request, a multi-step biological reasoning query that exceeded the framework’s analytical scope, and a large-dataset timeout where the system suggested a scalable alternative. The ablation data (§2.4) provide concrete examples of what each error layer catches in practice (Supplementary Table 6). At the schema boundary, Pydantic rejects hallucinated method names (e.g., `“deep learning”`) and case mismatches (e.g., `“RCTD”` vs. `“rctd”`), returning typed errors that list the valid options. Beyond the schema boundary, data-state checks catch missing prerequisites—for instance, requesting spatial visualization on an object lacking coordinates returns a `DataNotFoundError` identifying the missing field. Dependency checks catch a subtler case: a model may select a valid enumeration value whose package is not installed (e.g., `cell2location`), and the runtime returns a `DependencyError` naming available alternatives. The quantitative gap between these layers—98.3% validation under full schema vs. 20.4% without, and a further 3.8-percentage-point drop from validation to execution under bare schema (§2.4)—shows that each layer contributes non-redundantly to failure prevention.

These observations map onto a four-layer error architecture. (1) The *schema boundary* (Pydantic validation) catches invalid method names, type errors, and out-of-range values before any analysis code runs. (2) *Data-state checks* in tool wrappers catch missing spatial coordinates, absent preprocessing prerequisites, and invalid column references. (3) *Dependency checks* catch missing packages and provide installation guidance with available alternatives. (4) *Processing checks* catch algorithm convergence failures, numerical errors, and insufficient input data. Each layer converts the failure into a structured diagnostic response returned through the MCP tool interface, enabling correction or alternative analysis while preserving an auditable record.

These 31 scenarios exercise known edge cases; field deployment will inevitably surface additional failure modes, particularly in data-format variation and cross-platform coordinate conventions.

## 3 Discussion

The central finding of this work is that constraining an LLM’s output space to typed schemas and curated parameter fields substantially narrows variability in spatial transcriptomics workflows—without sacrificing the flexibility of natural-language interaction. The ablation traced a clear causal chain: typed constraints raised parameter validity from 20% to 98%, validity gated execution success, and descriptive guidance standardized method choice where multiple algorithms were available. These effects were largest for parameter-sensitive tasks and smallest where algorithms have robust defaults, consistent with a design whose value scales with the complexity of the parameter space. The case-study concordance experiment reinforces this: across four LLMs including the model used for the original case-study analyses, CARD deconvolution yielded identical outputs in every full-schema trial (*r* = 1.000, 780 cross-model pairs per sample), confirming that the published results are not contingent on a specific model. The DLPFC ground-truth benchmark extends this from consistency to external concordance: ChatSpatial achieved higher ARI against expert-annotated cortical layers than the two comparison systems (0.374 vs. 0.274 for SpatialAgent and 0.227 for STAgent), while maintaining perfect cross-replicate consistency.

Because each step is recorded as a structured tool call, reproducibility also enables composition: cross-method queries that would normally require separate scripting become single-prompt operations. In the case studies, a single prompt connected CellChat ligand–receptor results to Moran’s *I* spatial rankings (OSCC) and linked RCTD deconvolution outputs to CNV-cluster spatial patterning (HGSOC).

ChatSpatial occupies the high-constraint end of AI-assisted spatial analysis (Supplementary Table 4). STAgent [9] and SpatialAgent [10] provide complementary designs that emphasize broader automation through tool libraries, retrieval, templates, or generated code. The cross-system comparison (§2.4) quantifies the tradeoff: constrained tool selection achieved higher accuracy and consistency at the invocation level, and this advantage extended to full pipeline execution—but the benefit is specific to reproducible execution through a curated method set. Lower-constraint architectures may offer advantages for open-ended exploration or tasks outside a predefined repertoire. The same invocation layer also enables cross-ecosystem composition: R-based methods such as CellChat and RCTD coexist with Python-native methods such as infercnvpy. Learned models for semantic exploration, such as CellWhisperer, address a complementary problem and could operate as tools within the same orchestration framework.

GUI-based tools such as 10x Genomics Loupe Browser, Giotto Viewer, and web platforms such as Galaxy offer accessible alternatives for specific analytical tasks (Supplementary Table 5). ChatSpatial addresses a different use case: reproducible composition of specialized methods across Python and R. The approaches are complementary: GUI tools support interactive visualization and targeted exploration; typed-schema orchestration supports multi-method workflows that need explicit provenance across software ecosystems.

Several limitations define the current scope. Natural-language orchestration remains model-dependent: the prompt sensitivity experiment demonstrated 98.7% cross-paraphrase concordance under controlled conditions, but robustness to domain-specific jargon and ambiguous biological phrasing remains untested, and LLM updates can affect tool selection over time. The ablation and benchmark were tested on a limited number of datasets and LLMs; general reproducibility across diverse datasets, platforms, or upstream package versions remains to be established. ChatSpatial also inherits the limitations of the methods it orchestrates: tool selection reflects curatorial judgment, and wrapper maintenance is required as upstream packages evolve.

Deployability intersects with reproducibility in practice. The current command-line and MCP-client interface assumes familiarity with terminal workflows, which limits adoption among experimentalists who generate spatial data but rely on GUI-based tools for analysis. More broadly, dependence on external LLM APIs introduces latency, cost, and privacy considerations—particularly relevant for clinical datasets subject to institutional data-use agreements. Local model deployment can address privacy constraints but may alter tool selection behavior; the extent of this effect remains to be quantified.

Finally, the case-study validation relies on published analyses, so we cannot exclude the possibility that model priors influenced parameter selection; stronger evidence of generalizability will require blinded datasets and independent biological validation.

As spatial transcriptomics workflows continue to grow in analytical breadth, the gap between available methods and researchers’ ability to compose them reproducibly will widen. Schema-enforced orchestration is one architecture for closing that gap— constraining what the LLM can do while preserving the flexibility of natural-language interaction and keeping every analytical decision visible for review.

## 4 Methods

### 4.1 Schema Enforced Invocation

ChatSpatial is built on the Model Context Protocol (MCP) [11], an open protocol for exposing tools to LLM-enabled applications. The implementation uses a **schema-enforced execution model**: the LLM selects an MCP tool and fills typed parameter fields, while the analysis code is executed by versioned Python or R wrappers.

The 20 MCP tools (e.g., `deconvolve_data`, `identify_spatial_domains`) each accept a `method` parameter that routes to specific algorithm implementations. For instance, when a user requests deconvolution, the LLM cannot invent a method name; it must select from a pre-validated enumeration (e.g., `Literal[“rctd”, “card”, “cell2location”]`). The parameter-level breakdown is reported in §2.1.

By enforcing type safety and parameter validity before analysis code is executed, the architecture prevents syntax errors and invalid enum choices at the tool-call boundary, while leaving data quality, ambiguous intent, dependency issues, and upstream method limitations as explicit failure modes. For fixed tool choices and parameter values, execution follows deterministic software paths under fixed package and environment versions. Residual variability therefore resides mainly in intent-to-parameter mapping, which the schema constraints are designed to narrow. Our implementation supports the 2025-03-26 MCP specification through the Python SDK (v1.17.0+). Further details on the protocol architecture and system prompts are provided in the Supplementary Methods.

### 4.2 Method Integration

We integrated the full method library (Supplementary Table 2) through an MCP-based tool integration architecture designed to preserve analytical fidelity while presenting a consistent interface for the LLM. The architecture consists of three operational layers:

1. **MCP Tool Layer**. Each analysis function is registered as an MCP tool via the @mcp.tool() decorator, exposing it to the LLM with strict input/output schemas that enforce parameter validation and type safety.
2. **Method Dispatch Layer**. Registered tools perform data validation, coordinate management, and route requests to algorithm-specific implementations based on user-specified method parameters.
3. **Algorithm Wrapper Layer**. Method-specific wrappers handle data preparation (coordinate extraction, quality checks, format conversions), call the original software with appropriate parameters, and standardize outputs into consistent result objects.

For methods that are only available in R—such as CellChat for cell communication analysis—the wrapper layer utilizes `rpy2` to call R from within the Python-based server, automatically converting `AnnData` objects to `Seurat` objects and translating results back. This inter-language process is encapsulated within the wrapper and invisible to the user. A detailed example of this integration process for SpaGCN spatial domain identification is provided in the Supplementary Methods.

### 4.3 parameter Guidance

ChatSpatial embeds curated method guidance directly into MCP tool schemas. Each parameter field includes both machine-readable constraints and LLM-visible descriptions. For instance, in the spatial graph construction tool (`build_spatial_graph`), the description for `n_neighbors` encodes technology-specific heuristics: *“For 10x Visium (hexagonal grid), recommended k=6 for immediate neighbors; for single-cell resolution platforms like MERFISH, recommended k=10-15 based on cell density; for Xenium, consider Delaunay triangulation for natural cell neighborhoods*.*”* Similarly, the `clustering_resolution` description explains how lower and higher values change the granularity of detected tissue domains.

When the LLM processes a user request, it receives the relevant tool schema and can use these descriptions to propose parameter values within the allowed space. This is not automatic parameter optimization; it is documentation-guided parameter selection that remains visible to the researcher. The advantage is maintainability: method guidance can be updated by revising schema descriptions rather than retraining a model when new spatial technologies or best practices emerge. Further examples are provided in the Supplementary Methods.

### 4.4 Error Handling

ChatSpatial implements error handling designed for conversational interaction rather than silent recovery. The core design principle is that errors are converted into structured result objects and returned to the LLM, enabling it to observe failures and guide users toward solutions through natural language rather than requiring manual debugging. This is achieved through two key mechanisms: (1) a semantic exception hierarchy that categorizes errors by type (data issues, parameter errors, processing failures, dependency problems), enabling the LLM to provide targeted guidance; and (2) the @mcp_tool_error_handler decorator that wraps all MCP tools with type-aware error responses, maintaining protocol-level consistency while returning actionable diagnostic messages. Detailed implementation is provided in the Supplementary Methods.

### 4.5 Reproducibility Experiments

Two controlled experiments evaluated whether schema enforcement improves invocation consistency (ablation) and whether tool selection is robust to prompt variation (sensitivity). Both experiments shared an infrastructure: each LLM received a system prompt containing one of the schema variants, a fixed response-format instruction requiring JSON output ({`“tool_name”:…, “parameters”: {…`}}), and the user prompt. Responses were parsed, the tool name was resolved through a canonical alias map, and parameters were validated against the corresponding Pydantic model from ChatSpatial’s codebase—the same validation that gates real tool execution.

#### Schema-enforcement ablation

Three schema conditions were constructed from the production schema text. The *full schema* preserved the complete tool definition: parameter types, Literal enumerations, numeric bounds, defaults, and natural-language descriptions of method behavior and platform-specific recommendations. The *bare schema* retained types, enumerations, bounds, and defaults but removed all descriptive text, isolating the contribution of machine-readable constraints. The *no-schema* condition listed only tool names with one-sentence capability summaries, removing all typed parameter information. Eight prompts were selected to cover six of the eight analytical tools, spanning spatial domains, deconvolution, SVG detection, cell communication, spatial statistics, visualization, trajectory, and CNV analysis. Each prompt specified enough context for unambiguous tool selection (e.g., method hints, required reference identifiers) while leaving constrained parameters such as `method` for the model to infer from schema content. Three LLMs—Gemini 2.5 Flash, Claude Haiku 4.5, and GPT-5.4—were chosen as lightweight, cost-efficient representatives of three major providers; the case-study analyses used the more capable Claude Sonnet 4.5. All API calls used temperature *T* = 1.0 to maximize stochastic variation.

The invocation-level experiment (720 trials: 8 prompts *×*3 models *×*10 replicates *×*3 conditions) measured parse success, tool selection accuracy, and Pydantic validation rate. The end-to-end extension (270 trials: 3 tasks *×*3 models *×*10 replicates *×*3 conditions) executed validated invocations on an OSCC Visium dataset (GSE208253, Sample 1; 1,159 spots *×*15,215 genes), with each trial operating on an independent data copy to prevent cross-contamination.

Output concordance was computed as pairwise metrics within each condition *×* task cell: Adjusted Rand Index for cluster labels, Jaccard similarity of top-100 ranked genes, and Pearson correlation of cell-type proportion vectors. All reported confidence intervals are 95% percentile bootstrap intervals (10,000 resamples, seed = 42). Statistical comparisons across the three conditions used Kruskal–Wallis tests, with pairwise Mann–Whitney *U* tests and Bonferroni correction for post-hoc contrasts.

#### Prompt sensitivity

Five prompt groups were selected from the ablation prompt set (spatial domains, deconvolution, SVG detection, cell communication, Moran’s *I*). For each group, one canonical prompt was retained from the ablation and four semantically equivalent paraphrases were written, varying vocabulary and syntactic structure while preserving analytical intent and any required parameter hints (e.g., method name, reference identifiers). The experiment comprised 375 trials (5 groups *×* 5 paraphrases *×* 3 models *×* 5 replicates) under the full-schema condition at *T* = 1.0. Cross-paraphrase concordance was measured as the modal agreement of constrained parameters across paraphrases within each model group *×* cell, and compared with the within-paraphrase consistency from the ablation experiment.

#### Cross-system invocation comparison

To test whether schema enforcement confers advantages relative to other architectural designs, we extracted the full invocation context—tool catalog, system prompt, and routing instructions—from STAgent [9] (8 tools) and SpatialAgent [10] (72 tools). The same eight prompts were presented to the same three LLMs under each system’s context at *T* = 1.0, yielding 480 new trials (1,200 total across five conditions including the three ChatSpatial ablation conditions). Tool selection accuracy was evaluated by domain-expert annotation of whether the selected tool was appropriate for the prompted task.

#### End-to-end execution benchmark

To extend the comparison to full pipeline execution, we ran all three systems on a human lymph node Visium dataset (10x Genomics; 4,035 spots *×* 36,601 genes; not used in any case study) with identical natural-language prompts and the same LLM (Claude Sonnet 4, *T* = 1.0). Three tasks were tested—spatial domain identification, spatially variable gene detection, and cell–cell communication (ligand–receptor analysis)—with 5 replicates per system per task (45 total trials). Each system was invoked through its native API: ChatSpatial via direct tool execution after LLM-based parameter selection, STAgent via its LangGraph agent loop (`invoke_our_graph`), and SpatialAgent via its Plan-Act-Conclude agent (`SpatialAgent.run`). Each trial ran in an isolated environment with a 10-minute timeout. Output concordance was computed as pairwise Adjusted Rand Index (spatial domains), Jaccard@100 (SVG rankings), and Jaccard@50 (ligand–receptor pair overlap) across successful replicates within each system; for spatial domains, ARI was computed on the intersection of common spot barcodes to handle minor preprocessing differences in spot filtering across systems.

#### Case-study workflow concordance

To bridge the ablation findings to the case-study results, we repeated the CARD deconvolution step from the OSCC workflow (§2.3, Step 2) under full-schema and no-schema conditions. The prompt matched the case-study analytical intent: “Deconvolve cell types in this OSCC sample using CARD with the HNSCC single-cell reference from Puram et al. (reference_data_id=‘puram_ref’, cell_type_key=‘cell_type’).” Four LLMs were tested—Gemini 2.5 Flash, Claude Haiku 4.5, GPT-5.4, and Claude Sonnet 4.5—on Samples 1 and 9 (160 total trials: 2 samples × 2 conditions × 4 models × 10 replicates, *T* = 1.0). Output concordance was measured by pairwise Pearson *r* of flattened proportion matrices with bootstrap 95% CIs (10,000 resamples). CARD was selected for this bridge experiment because it is deterministic: any output variation is attributable entirely to schema-condition-induced parameter variation.

#### Ground-truth benchmark

To evaluate output correctness against independent expert annotations, we tested all three systems on the DLPFC Visium dataset [27], which provides expert-annotated cortical layer labels (L1–L6 and white matter) for 12 samples across three donors. Three samples were selected (151673, 151507, 151669; one per donor) via the `spatialLIBD` R/Bioconductor package. Each sample was preprocessed (normalization, HVG selection, PCA, spatial neighbors) and converted to h5ad format; ground-truth labels were stored separately and excluded from the data provided to systems to prevent information leakage. An open-ended prompt was used: “This is human dorsolateral prefrontal cortex (DLPFC) Visium data. Identify spatial domains in this dataset.” Each system ran 10 replicates per sample (90 total trials; Claude Sonnet 4, *T* = 1.0, 15-minute timeout). Ground-truth concordance was measured by ARI and NMI computed on the intersection of spots with both predicted labels and expert annotations. Cross-replicate concordance was measured by pairwise ARI as in the end-to-end benchmark. All 95% CIs were computed using hierarchical bootstrap (resampling samples with replacement, then replicates within each resampled sample; 10,000 iterations) to account for the nested trial structure (*n* = 3 independent samples). All prompts, schema variants, analysis scripts, and aggregate result tables are provided in the reproducibility repository. Raw provider response checkpoints can be regenerated by the API-dependent scripts and are not required for inspecting the committed numerical results.

## Acknowledgements

We thank the MCP development team at Anthropic for creating the framework that enables ChatSpatial’s LLM-native architecture. We are grateful to Dr. Jade Wang (Department of Statistics, Texas A&M University) for valuable feedback and insightful suggestions on the design of the case study analyses. We thank Jiahao Mai (Southern Medical University) for assistance with figure design and visualization improvements.

## Supplementary Information

Supplementary information is available as a separate PDF. Supplementary Tables 1–4 are provided as standalone table files, and Supplementary Tables 5–6 are included in the Supplementary Information PDF.

## Data availability

All public datasets used for benchmarking and validation in this study are listed below and documented in a dedicated reproducibility repository (https://github.com/cafferychen777/ChatSpatial-Reproducibility):

- Oral Squamous Cell Carcinoma (OSCC) Visium dataset: GEO accession GSE208253 [21]
- High-Grade Serous Ovarian Carcinoma (HGSOC) Visium dataset: GEO accession GSE211956 [22]
- HNSCC single-cell reference: GEO accession GSE103322 [20]

The end-to-end execution benchmark (Section 2.4) used a human lymph node Visium dataset from the 10x Genomics public dataset collection. The ground-truth benchmark used three DLPFC Visium samples (151673, 151507, 151669) from the `spatialLIBD` R/Bioconductor package [27]. Additional datasets used for functional validation (Sections 2.2 and 2.5) include SPOTS (GSE198353), Visium multi-sample benchmarks (GSE254652, GSE243275), MERFISH (GSE113576), seq-FISH (GSE133244), STARmap (https://www.wangxiaolab.org/data-portal-1), Slide-seq (Broad Institute Single Cell Portal SCP354), and Xenium (10x Genomics public dataset). All datasets are publicly available, and no custom datasets were generated for this study.

## Code availability

The source code for ChatSpatial (v1.2.10) is publicly available on GitHub under the MIT license at https://github.com/cafferychen777/ChatSpatial. A pre-built Docker image is available at ghcr.io/cafferychen777/chatspatial:v1.2.10. Comprehensive documentation is hosted at https://docs.cafferyang.com/. All analysis scripts used to generate the figures and results in this paper are available in the reproducibility repository at https://github.com/cafferychen777/ChatSpatial-Reproducibility.

Installation: `uv pip install chatspatial` (requires the `uv` package manager; https://docs.astral.sh/uv/). The base installation supports the core MCP server and Python-native analysis wrappers without pulling compiled optional packages such as `gseapy`, R packages, or deep-learning dependencies. Users who need R-based methods, enrichment analysis, or the complete method catalog can install the advanced `chatspatial[full]` extra, whose documented platform prerequisites include Rust/LLVM on Intel macOS when upstream wheels are unavailable. A pre-built Docker image (ghcr.io/cafferychen777/chatspatial:v1.2.10) provides a validated full-stack environment with dependencies pre-resolved.

## Declarations

### Ethics approval and consent to participate

Not applicable. This study only used publicly available datasets and did not involve human participants or animal experiments.

### Consent for publication

Not applicable.

### Author contributions

C.Y.: Conceptualization, Methodology, Software, Validation, Formal Analysis, Investigation, Data Curation, Writing – Original Draft, Visualization. X.Z.: Conceptualization, Methodology, Resources, Writing – Review & Editing, Supervision, Project Administration, Funding Acquisition. J.C.: Conceptualization, Resources, Writing – Review & Editing, Supervision, Project Administration, Funding Acquisition. All authors read and approved the final manuscript.

